# RNAStabFormer: Region-Aware Multi-Task Hybrid Learning for RNA Stability Prediction from Pulse-Chase Transcriptomics

**DOI:** 10.64898/2026.06.19.733389

**Authors:** Shuo Wang, Chuqiao Zhang

## Abstract

RNA stability is a major post-transcriptional regulator of gene expression, yet sequence-based prediction from pulse-chase transcriptomics remains difficult because labels depend on time window, quantification region, and replicate quality. We present RNAStabFormer, a controlled RNA stability framework centered on a Region-aware Multi-task Hybrid Transformer (RAMHT). RAMHT encodes 5′UTR, CDS, and 3′UTR nucleotide context, adds a CDS codon stream, upgrades engineered sequence features into a tabular interaction branch, and uses gated multi-task regression to predict four ENCODE BrU-seq/BruChase-seq stability proxies, with exon total 6 h/0 h as the primary task. Across 26 outer splits including 23 chromosome holdouts, a heterogeneous three-member RAMHT ensemble achieves 0.773 mean Pearson correlation on the primary task, statistically matching an engineered-feature XGBoost baseline (0.773; mean paired delta +0.000004; bootstrap 95% CI [− 0.003845, +0.004077]; Wilcoxon *p* = 0.8613). The ensemble improves over the strongest single RAMHT member (0.768 to 0.773; 23/26 split wins), while a strict nested XGBoost+RAMHT blend reaches 0.775. Same-split checks also exceed frozen full-length mRNA-LM embeddings (0.760) and public LAMAR-DR transfer (0.180). Gate, ablation, and recoding analyses show that engineered sequence grammar remains dominant, while nucleotide and codon branches provide complementary CDS-local signal. RNAStabFormer narrows the gap between neural RNA sequence modeling and strong tabular baselines while retaining an extensible architecture for interpretation and biological integration.

## I. Introduction

RNA stability determines how long a transcript remains available for regulatory or translational activity after it is synthesized. It is therefore a central component of post-transcriptional gene regulation, complementary to transcription rate, splicing, export, translation, and decay. Stability is also a sequence-mediated phenotype: untranslated regions, coding sequence, codon usage, local nucleotide grammar, RNA structure, and binding motifs can all contribute to transcript retention and degradation. This makes RNA stability an appealing target for sequence models, but also a challenging benchmark for machine learning.

Pulse-chase transcriptomic assays such as BrU-seq and BruChase-seq measure nascent and retained RNA after metabolic labeling [1], [2]. A supervised stability target is not observed directly; it is derived from ratios between chase-time signals after choosing a quantification region, filtering low-signal denominators, reconciling biological replicates, and aggregating across cellular contexts. Small choices in this construction can change the apparent predictability of the task. For this reason, a convincing model must be evaluated with fixed cohorts, strict train/validation/test separation, and strong engineered-feature baselines.

Recent neural and language-model approaches show that mRNA sequence contains predictive information about degradation rates [3]–[5], but reported scores are tied to different labels: Saluki uses mammalian consensus half-life labels, mRNA-LM uses full-length mRNA property benchmarks, and LAMAR-DR is tuned on 3′UTR reporter half-lives. Our labels are endogenous pulse-chase retention proxies with fixed chromosome holdouts, so we evaluate external models under the same outer-test protocol whenever possible. In this setting, XGBoost [6] trained on engineered sequence features remains the strongest non-neural stress test. The key question is whether a hybrid neural model can match strong tabular and pretrained-LM alternatives while retaining sequence-aware extensibility.

We address this question with RNAStabFormer, a framework centered on RAMHT. The model follows three design principles. First, transcript regions should be encoded separately before being fused, because 5′UTR, CDS, and 3′UTR have different sequence grammars and biological roles. Second, CDS should be represented at both nucleotide and codon resolutions. Third, engineered sequence grammar should be treated as a first-class input rather than a weak auxiliary covariate, because it captures high-signal k-mer and composition patterns that are difficult to learn from approximately ten thousand genes.

We make four contributions: strict ENCODE BrU-seq/BruChase-seq stability labels [7], [8] using GENCODE annotation [9]; RAMHT with region-aware nucleotide encoding, a codon branch, tabular interaction encoding, gated fusion, and multi-task heads; a 26-split evaluation against XGBoost, mRNA-LM frozen embeddings, LAMAR-DR transfer, and local deep sequence baselines; and gate, residual, ablation, feature-importance, and synonymous-recoding analyses that clarify the captured signal.

## II. Data and Task Construction

### A. Pulse-Chase Stability Proxies

We use ENCODE BrU-seq/BruChase-seq signal tables generated by the Ljungman laboratory as part of the ENCODE deeply profiled cell-line resource [7], [8]. The data cover 16 cell lines, three chase times (0 h, 2 h, and 6 h), and two biological replicates per cell line. We process two signal channels: gene-sense quantification and exon-sense quantification. These channels produce related but distinct views of retained RNA, with exon-sense total-chase signal forming the primary prediction task in this work.

For gene *g*, cell line *c*, replicate *r*, and channel-specific signal *s*_*t,gcr*_, we define two retention ratios:

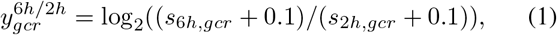

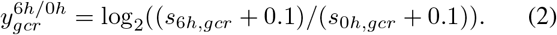

All labels require *s*_0*h*_ ≥ 0.5; 6 h/2 h labels additionally require *s*_2*h*_ ≥ 0.5. Gene-cell measurements are excluded when paired biological replicates differ by more than 1.0 log2 unit. Final gene-level labels are pass-only cross-cell consensus values over genes observed in at least eight cell lines. Table I summarizes the resulting cohorts.

**TABLE I.** RNA Stability Proxy Labels.

| Label | Consensus | Model genes | QC pass | Role |
| --- | --- | --- | --- | --- |
| Gene 6 h/2 h | 8428 | 8428 | 0.836 | auxiliary |
| Gene 6 h/0 h | 9848 | 9848 | 0.839 | auxiliary |
| Exon 6 h/2 h | 9765 | 9018 | 0.849 | auxiliary |
| Exon 6 h/0 h | 10678 | 9881 | 0.836 | primary |

### B. Transcript Inputs

GENCODE v29 is used to assign representative transcripts and extract 5′UTR, CDS, and 3′UTR sequences. Each gene is represented by raw regional nucleotide strings, a CDS codon string, and 1336 engineered sequence features. Engineered features include regional lengths, nucleotide composition, GC/AU fractions, 3-mer and 4-mer frequencies, codon summaries, and selected regulatory motif counts. Gene identifiers, chromosome, strand, biotype, and quality-control metadata are not used as predictors.

## III. RNAStabFormer

### A. Region-Aware Hybrid Architecture

Figure 1 shows the RAMHT architecture, the core model inside RNAStabFormer. The model uses independent nucleotide encoders for 5′UTR, CDS, and 3′UTR. Each encoder contains nucleotide embeddings, positional embeddings, convolutional motif stems in selected variants, Transformer blocks [10], and attention pooling. A separate CDS codon branch tokenizes coding sequence into codon categories and applies a codon encoder before projection into the shared latent dimension.

**Fig. 1.**
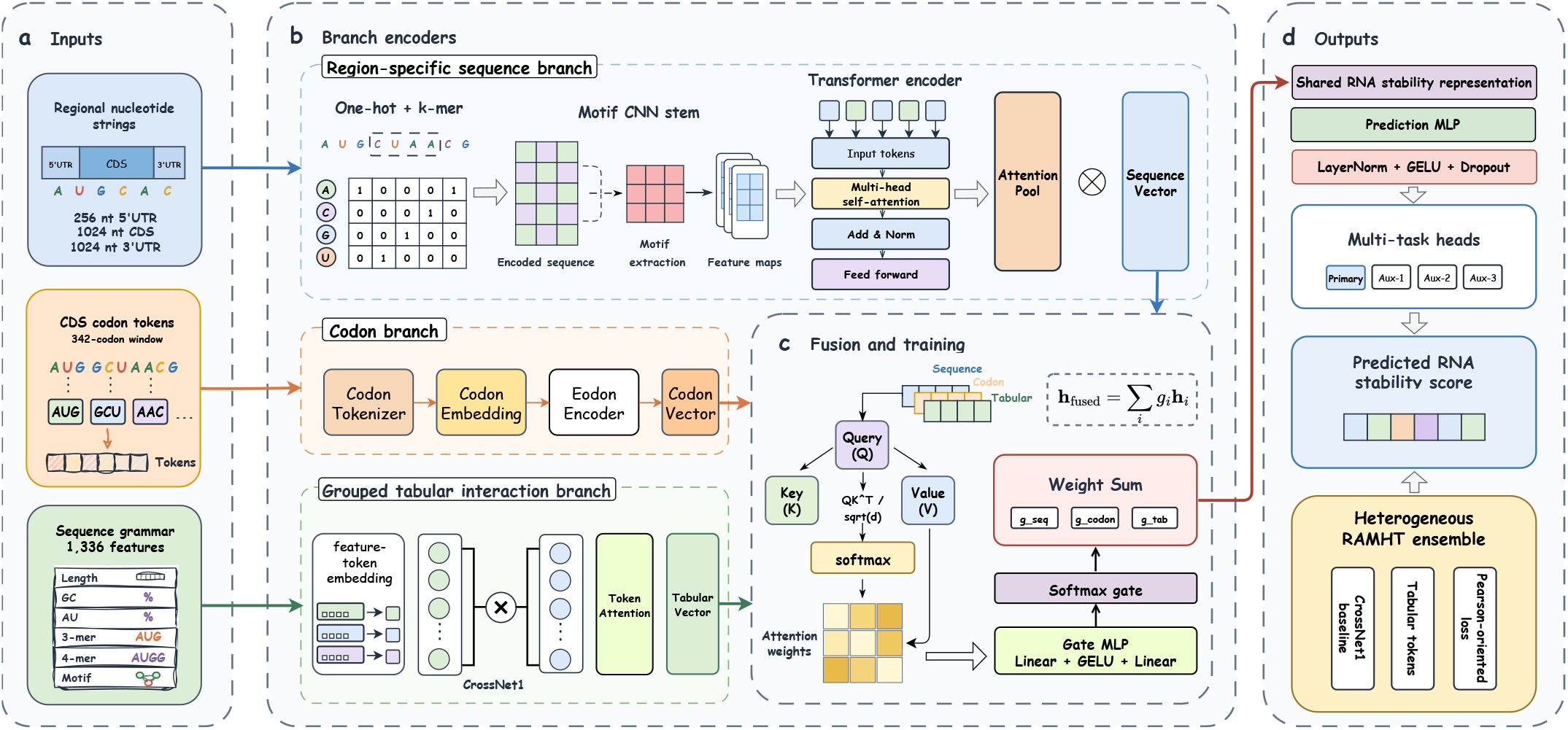
RAMHT as the core model architecture of RNAStabFormer. (a) Inputs include regional nucleotide strings, CDS codon tokens, and 1336 engineered sequence-grammar features. (b) Branch encoders convert these views into sequence, codon, and tabular vectors using motif-CNN/Transformer attention, codon encoding, grouped feature tokens, and CrossNet1-style interactions. (c) QKV attention and a softmax Gate MLP fuse the three branch vectors into **h**_fused_. (d) A shared prediction MLP, four task heads, and a heterogeneous RAMHT ensemble output RNA stability scores.

**Fig. 2.**
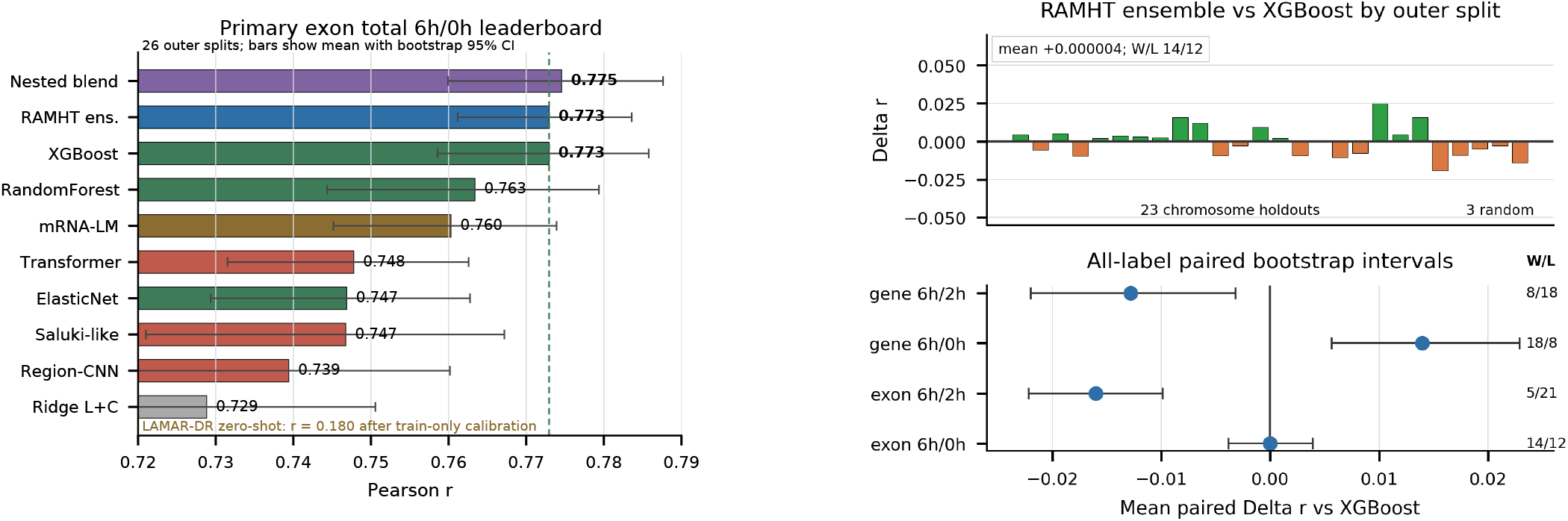
Primary leaderboard and paired RAMHT-vs-XGBoost deltas. Left: 26-split Pearson means with bootstrap 95% CIs. Right: primary split-level deltas and all-label paired bootstrap intervals with wins/losses.

**Fig. 3.**
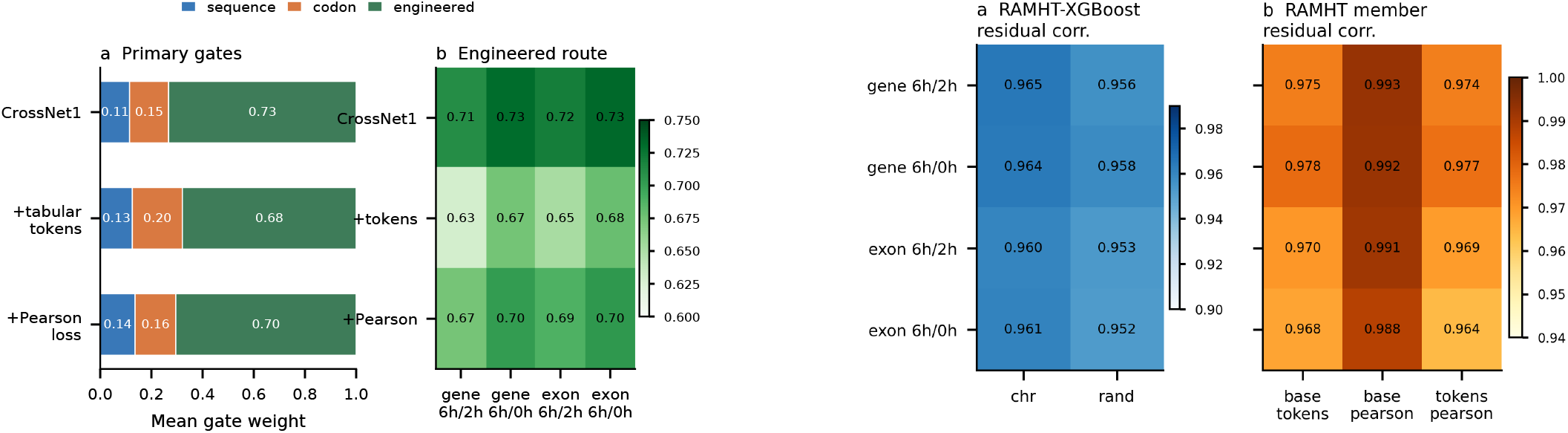
Gate allocation and residual complementarity. Engineered features dominate gate weights while sequence/codon branches remain nonzero; residual heatmaps show high RAMHT–XGBoost and within-RAMHT member error similarity.

Within the sequence, tabular-token, and fusion blocks, token interactions are modeled with scaled dot-product attention:

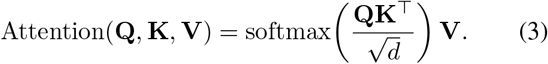

The engineered-feature branch is designed as an interaction module rather than a plain vector append. In the strongest tabular-token variant, features are grouped into biologically interpretable blocks, embedded as feature tokens, and allowed to interact with sequence and codon summaries through attention. Other ensemble members use a CrossNet-style tabular interaction block [11], [12] or a correlation-oriented training objective. A learned gate then combines the nucleotide, codon, and engineered-feature streams. The fused representation is passed to task-specific regression heads for the four labels.

Let **h**_seq_, **h**_codon_, and **h**_tab_ denote the sequence, codon, and tabular branch vectors, respectively. RAMHT computes a sample-specific softmax gate and forms the shared stability representation as

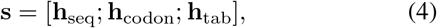

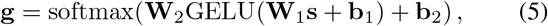

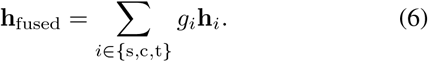

Each task head then predicts

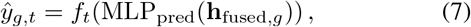

where *t* indexes the primary or auxiliary stability label.

### B. Heterogeneous RAMHT Ensemble

The final RAMHT predictor is an equal-weight ensemble of three compatible members trained on identical outer splits. The members are: a CrossNet1 hybrid RAMHT baseline, a CrossNet1 model with tabular feature-token attention, and a CrossNet1 model with a Pearson-oriented loss term. The ensemble intentionally combines small architectural and objective-level differences rather than multiple random seeds of the same model. This design targets variance reduction and modest complementarity while preserving a single conceptual architecture.

When XGBoost distillation is used during training, teacher predictions are generated out-of-fold inside the outer-training data only. Thus, the neural student never receives in-sample teacher predictions for outer-test genes, and teacher information cannot leak across the final evaluation boundary.

### C. Leakage-Controlled Multi-Task Splits

For each outer split, genes that are test genes for any label are excluded from training and validation for all tasks. Genes that are validation genes for any label are excluded from training unless they are already in the union test set. This conservative rule prevents a shared encoder from observing a gene in one task and being evaluated on the same gene in another task. Feature standardization, target normalization, early stopping, and weight selection are performed using training or validation data only.

## IV. Experimental Design

### A. Baselines

The main same-split comparator is XGBoost trained on the same engineered sequence features as the RAMHT tabular branch. XGBoost is an intentionally strong baseline because it captures nonlinear k-mer, composition, and regional feature interactions efficiently in small-to-medium tabular datasets. We also evaluate ElasticNet [13], RandomForest [14], raw-sequence deep networks, a Saluki-like CNN-GRU model, weak ridge baselines, and earlier RAMHT variants. The paper’s primary claim is made against XGBoost because it is the strongest deployed baseline for the primary label.

To benchmark external RNA language models fairly, we adapt public checkpoints to the same 26 outer splits instead of comparing incompatible literature numbers. For mRNA-LM [4], the official 5′UTR, CDS, and 3′UTR BERT checkpoints are frozen, each gene is represented by the concatenated regional mean embedding, ridge regularization is selected using outer-validation genes, and the final head is trained only on outer-training genes. For LAMAR-DR [5], public 3′UTR degradation scores are linearly calibrated on outer-training genes and applied to outer-test genes. These transfer baselines test whether recent RNA foundation models solve the present endogenous pulse-chase label under the same evaluation boundary as RAMHT.

### B. Training and Implementation Details

Deep models are trained with masked multi-task regression [15], so a gene contributes only to labels that are available for that gene. Continuous targets are normalized using training genes only and transformed back for reporting. The main neural objective uses robust regression loss [16] with early stopping on outer-validation performance; selected variants add a Pearson-oriented loss term to directly optimize ranking agreement. For an outer-training set *D*, the objective can be written as

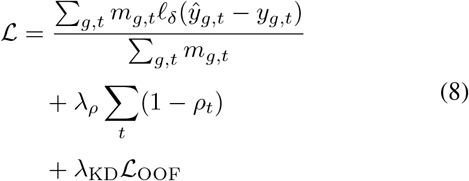

where *m*_*g,t*_ masks unavailable labels, ℓ_δ_ is the Huber loss, *ρ*_*t*_ is the validation-oriented Pearson correlation term for task *t*, and _OOF_ is used only when out-of-fold teacher distillation is enabled. Adam-family optimization [17], gradient clipping, and validation-based checkpoint selection are used for all neural models. The default wide-window configuration encodes 256 nt of 5′UTR, 1024 nt of CDS, 1024 nt of 3′UTR, and 342 CDS codons; memory-safe screening variants use shorter but proportional windows.

Model development used small chromosome screens to compare candidate architectures. The final three-member ensemble was then evaluated on the full 26-split protocol. All reported paired tests are computed on the locked outer-test predictions from this final evaluation.

### C. Outer Splits and Metrics

The evaluation contains 26 outer splits: three repeated-random splits and 23 chromosome-holdout splits. Chromo-some holdout is emphasized because it tests generalization to genes from held-out chromosomes rather than random genes from the same genomic distribution. Pearson correlation is the primary metric, with Spearman correlation and *R*^2^ retained as supporting diagnostics. All comparisons are paired at the split level. For the primary RAMHT-vs-XGBoost comparison, we report paired differences, wins/losses, paired bootstrap 95% confidence intervals, Wilcoxon signed-rank tests, and paired sign-permutation tests.

## V. Results

### A. RAMHT Matches XGBoost on the Primary Stability Task

The strongest RAMHT ensemble matches XGBoost on the exon total 6 h/0 h primary label across the full 26-split evaluation (Table II). Mean Pearson correlation is 0.772953 for the RAMHT ensemble and 0.772948 for XGBoost. Chromosome-holdout means are also nearly identical: 0.7732 for the RAMHT ensemble and 0.7723 for XGBoost. A strict nested XGBoost+RAMHT blend obtains the highest mean value (0.774567), but it is reported as a hybrid system rather than as standalone RAMHT.

**TABLE II.** Primary Exon Total 6 h/0 h Same-Split Leaderboard.

| Model | 26-split $r$ | Chr. $r$ | Random $r$ |
| --- | --- | --- | --- |
| Train mean | 0.000000 | 0.000000 | 0.000000 |
| LAMAR-DR zero-shot | 0.180130 | 0.181167 | 0.172179 |
| Ridge length+composition | 0.728836 | 0.728135 | 0.734212 |
| Saluki-like | 0.746756 | 0.746378 | 0.749650 |
| Transformer | 0.747786 | 0.747195 | 0.752312 |
| RandomForest | 0.763400 | 0.761903 | 0.774875 |
| mRNA-LM frozen ridge | 0.760312 | 0.759638 | 0.765474 |
| XGBoost | 0.772948 | 0.772275 | 0.778110 |
| 3-member RAMHT ensemble | 0.772953 | 0.773232 | 0.770809 |
| Nested XGBoost+RAMHT | 0.774567 | 0.773965 | 0.779182 |

**TABLE III.** Paired Primary-Task Tests Over 26 Splits.

| Comparison | Mean $\Delta r$ | W/L | Wilcoxon $p$ |
| --- | --- | --- | --- |
| Ens. vs. XGBoost | +0.000004 | 14/12 | 0.8613 |
| Ens. vs. CrossNet1 | +0.005374 | 23/3 | $7.45 \times 10^{-7}$ |
| mRNA-LM vs. XGBoost | -0.012637 | 4/22 | $1.91 \times 10^{-5}$ |
| Blend vs. XGBoost | +0.001619 | 18/8 | 0.0463 |

**TABLE IV.** Chromosome-Holdout Means Across Stability Proxies.

| Label | XGBoost | RAMHT ens. | Delta |
| --- | --- | --- | --- |
| Gene 6 h/2 h | 0.4975 | 0.4871 | -0.0104 |
| Gene 6 h/0 h | 0.5357 | 0.5518 | +0.0161 |
| Exon 6 h/2 h | 0.5573 | 0.5435 | -0.0138 |
| Exon 6 h/0 h | 0.7723 | 0.7732 | +0.0010 |

The external transfer checks strengthen the same-split claim. A frozen full-length mRNA-LM ridge baseline reaches 0.760 mean Pearson, outperforming local raw-sequence neural baselines but remaining below the RAMHT ensemble by 0.0126 Pearson on average. LAMAR-DR is a strong RNA foundation model in its own 3′UTR reporter-degradation setting, but its public checkpoint reaches only 0.180 mean Pearson after train-only calibration on our primary pulse-chase label. Together, these results show that RAMHT is competitive not only with local deep baselines but also with external pretrained RNA representations under the same outer-test protocol.

Paired tests indicate a match rather than a significant win over XGBoost. The mean split-level Pearson delta is +0.000004, the median delta is +0.001089, the ensemble wins 14 splits and loses 12, the paired bootstrap 95% interval is [− 0.003845, +0.004077], Wilcoxon *p* = 0.8613, and paired permutation *p* = 0.9985. Thus, RAMHT removes the practical performance gap to XGBoost on the primary task but does not yet prove significant superiority. The nested XGBoost+RAMHT blend gives the best mean performance and nominal Wilcoxon *p* = 0.0463, so we treat it as the strongest operational system.

### B. Ensembling Substantially Improves Single RAMHT Members

Although the RAMHT ensemble does not significantly exceed XGBoost, it clearly improves over the strongest single RAMHT member. The baseline CrossNet1 RAMHT reaches 0.767579 mean Pearson on the primary label, whereas the three-member ensemble reaches 0.772953. The ensemble wins 23 of 26 paired splits against CrossNet1, with a paired bootstrap interval fully above zero ([+0.003759, +0.007086]), Wilcoxon *p* = 7.45 *×* 10^−7^, and paired permutation *p* = 5.0 *×* 10^−5^. This result indicates that the performance gain is not a single-split fluctuation and that heterogeneous RAMHT members capture useful model-level diversity.

The tabular-token member is especially informative. It is the strongest single member on chromosome holdout (0.772559), nearly matching XGBoost by itself. This suggests that upgrading the engineered-feature branch from a simple MLP to a feature-token interaction module is one of the most effective architectural changes for the current data regime.

### C. Auxiliary Labels Reveal Where the Hybrid Model Helps

The auxiliary labels provide an important boundary around the primary-task claim. Across all four labels, the RAMHT ensemble is slightly below XGBoost on average (mean paired delta − 0.003720), with a bootstrap interval crossing zero ([− 0.008005, +0.000590]). Gene total 6 h/0 h is the most favorable auxiliary label for RAMHT: the ensemble reaches 0.5518 chromosome-holdout mean Pearson compared with 0.5357 for XGBoost. Gene late and exon late remain more difficult, and XGBoost remains stronger on those endpoints. Thus, the present evidence supports a high-performance primary RAMHT model and selective auxiliary gains, not universal dominance over XGBoost.

### D. Gates and Residuals Explain the Current Performance Boundary

Gate summaries show that engineered sequence grammar remains the dominant information route in hybrid RAMHT. On the primary label, the standard CrossNet1 RAMHT allocates mean gate weights of 0.115 to nucleotide sequence, 0.153 to codons, and 0.733 to engineered features. The tabular-token variant allocates 0.126, 0.195, and 0.679, respectively, increasing codon contribution while keeping engineered features dominant. The Pearson-loss variant shows a similar pattern, with 0.136, 0.159, and 0.705.

Residual correlations further clarify why the ensemble matches but does not significantly beat XGBoost. Primary-task residuals between the RAMHT ensemble and XGBoost have mean correlation 0.960 and median correlation 0.962. Residual correlations among RAMHT members are also high (mean 0.973). The models are therefore complementary enough for stable ensembling, but their errors are far from orthogonal. This explains the near-zero paired delta against XGBoost despite clear gains over single RAMHT members.

### E. Sequence Analyses Point to CDS-Local Grammar

Additional ablations support a biologically plausible sequence signal rather than a purely statistical artifact. In deep raw-sequence ablations, removing CDS causes the largest mean chromosome-holdout loss, about 0.05 Pearson, whereas removing 5′UTR or 3′UTR has a smaller effect. CDS-only input retains most raw-sequence performance, suggesting that coding-region sequence grammar is a major carrier of the supervised signal.

Engineered-feature screens agree with this region-level pattern. Group permutation importance ranks 4-mer and 3-mer features highest, with mean Pearson drops of 0.055 and 0.052, respectively, followed by composition and length features. Cross-label feature associations are dominated by GC-rich regional k-mers and composition features, while a small AU-rich motif panel is informative but insufficient by itself.

Thus, the predictive grammar is distributed across many local sequence features rather than explained by a single canonical motif.

The strongest mechanistic hint comes from synonymous CDS recoding. GC-min and GC-max synonymous recoding preserve amino-acid sequence but change synonymous codon choice and local nucleotide grammar; both perturbations shift model predictions, including large shifts on the primary exon total label. These findings are compatible with codon-usage-sensitive decay, translation-coupled stability, coding-region RBP binding, or surveillance-related sequence patterns. They remain in-silico association evidence, however, and should be treated as hypotheses for future perturbation or reporter assays rather than direct causal validation.

## VI. Discussion

RNAStabFormer shows that a hybrid neural model can reach the performance level of a strong engineered-feature XGBoost baseline and exceed a fair frozen mRNA-LM full-length embedding baseline on the primary RNA stability task. This is a hard comparison in the current data regime: XGBoost captures nonlinear engineered-feature interactions, whereas mRNA-LM contributes pretrained full-transcript representations. The final RAMHT ensemble closes this gap without using chromosome, gene identity, or quality-control metadata.

The design evidence supports a hybrid direction for RNA stability modeling. The engineered-feature branch receives the largest gate weight, confirming that k-mer, composition, and codon summaries remain highly informative at this sample size. Yet nucleotide and codon branches retain nonzero gate weights, heterogeneous ensembling improves over single RAMHT members, and the nested XGBoost+RAMHT blend gives the best mean primary performance. Thus, RAMHT turns a strong tabular predictor into a sequence-aware framework rather than a plain raw-sequence replacement.

The external language-model checks reinforce the need for matched evaluation: mRNA-LM frozen embeddings are competitive but remain below RAMHT, and LAMAR-DR transfer is weak for this endogenous pulse-chase label. The framework can also absorb longer context, RNA-binding protein tracks, miRNA seed matches, structure predictions, cell-line covariates, or pretrained RNA embeddings while preserving leakage-controlled evaluation. The current benchmark therefore establishes a high-performance, interpretable RAMHT ensemble and a clear path for future biological validation.

This modularity also makes RAMHT useful as a research platform rather than only a leaderboard model. Feature-token attention can be audited to identify which engineered sequence groups dominate individual predictions, gate weights can be stratified by transcript class or stability regime, and synonymous recoding tests can be extended from in-silico perturbation to reporter assays. These analyses would help separate general sequence grammar from cell-state specific regulation and could guide targeted additions such as RBP occupancy, miRNA seed context, RNA structure accessibility, or condition-specific decay factors [18]–[20].

## VII. Conclusion

We introduced RNAStabFormer, a region-aware multi-task hybrid framework for RNA stability prediction from pulse-chase transcriptomics. Its core model, RAMHT, integrates regional nucleotide encoders, a CDS codon stream, engineered-feature interaction modules, gated fusion, and task-specific heads. On the primary exon total 6 h/0 h label, a heterogeneous three-member RAMHT ensemble achieves 0.773 mean Pearson correlation across 26 outer splits, matching XGBoost, exceeding frozen mRNA-LM and LAMAR-DR transfer, and significantly improving over single RAMHT members. Mechanistic sequence analyses further support CDS-local, codon-sensitive hypotheses, making RAMHT a high-performance and extensible neural framework for RNA stability prediction.

